# Genomic signatures of positive selection and relaxed selective constraints associated with convergent dietary specialisation

**DOI:** 10.64898/2026.11.05.764531

**Authors:** Marta Olivé-Muñiz, Vadim A. Pisarenco, Nuria Macías-Hernández, Cristina Frías-López, Miquel A. Arnedo, Alejandro Sánchez-Gracia, Sara Guirao-Rico, Julio Rozas

**Affiliations:** Departament de Genètica, Microbiologia i Estadística, Universitat de Barcelona, Barcelona, Spain; Institut de Recerca de la Biodiversitat (IRBio), Universitat de Barcelona, Barcelona, Spain; Department of Animal Biology, Edaphology and Geology, University of La Laguna, Tenerife, Canary Islands, Spain; Departament de Biologia Evolutiva, Ecologia i Ciències Ambientals, Universitat de Barcelona, Barcelona, Spain

**Keywords:** Adaptive radiation, diet specialisation, convergent evolution, comparative genomics, heavy metals, spiders

## Abstract

Island radiations provide ideal frameworks for studying the molecular basis of convergent evolution. The rapid diversification of *Dysdera* spiders in Canary Islands provides a striking example, with repeated dietary shifts towards specialisation on woodlice, accompanied by morphological, metabolic, and behavioural modifications that facilitate the exploitation of this heavy-metal-rich prey. Here, we investigate two independent events of dietary specialisation on different islands. By analysing six high-quality genome-assemblies, including three newly generated in this study, we investigate the genomic signals of positive selection and relaxed selection in protein-coding genes, upstream regulatory regions, and gene family turnover rates.

We detected signatures of positive selection associated with the two specialists, accompanied by a twofold increase in gene loss rates relative to generalist ancestors. Notably, positive selection targeted a smaller, functionally coherent set of genes, potentially involved in detoxification likely through heavy metal excretion. In contrast, relaxation of selection affected a broader set of genes associated with functions that may have become dispensable under the new dietary regime. Furthermore, we found contractions in gene families associated with vesicular transport, potentially reflecting a reduced reliance on vesicle-mediated metal accumulation. Overall, our findings suggest that similar ecological pressures can drive parallel, yet non-identical, adaptive and non-adaptive responses, pointing to partially predictable evolutionary trajectories in an island radiation.

## INTRODUCTION

Oceanic archipelagos are classic model systems for studying a wide range of evolutionary processes (Emerson, 2002; Losos & Ricklefs, 2009; Yoder et al., 2010). Their isolation, well-defined boundaries, and known geological histories, facilitate the study of repeated evolution in short time periods. As these insular systems are often geologically young, ecological niches are typically rapidly filled through the diversification of a few early colonisers, resulting in taxa with recent shared ancestry, genomic backgrounds and common developmental constraints (Elmer & Meyer, 2011; Losos & Ricklefs, 2009). These features increase the likelihood of detecting genomic signatures of rapid diversification, rather than them being eroded by subsequent evolutionary processes (Cerca, Cotoras, Bieker, et al., 2023; Elmer & Meyer, 2011).

Dietary specialisation is a well-known case of convergent evolution that has long intrigued evolutionary biologists (Day et al., 2016; Futuyma & Moreno, 1988; Liem, 1980). Since food selection and trophic resource partitioning have important evolutionary and ecological consequences (Slatyer et al., 2013; Wilson et al., 2008), they often act as main drivers of adaptive radiation. In oceanic islands, low levels of interspecific competition during early colonisation can promote rapid population growth, leading to density compensation, whereby island populations reach higher densities than their mainland counterparts (Yoder et al., 2010). This increase is often followed by an intensification of intraspecific competition, which can promote niche partitioning and speciation (MacArthur et al., 1972; Svanbäck & Bolnick, 2005; Yoder et al., 2010), fostering the emergence of diverse dietary strategies. When these strategies evolve independently under comparable selective regimes rather than shared ancestry, they are clear candidates to represent adaptive convergence (Arendt & Reznick, 2008; Losos, 2011; Losos et al., 1998; Muschick et al., 2012; Roycroft et al., 2021; Yue et al., 2024).

Diet specialisation can be adaptive if it reduces the competition costs or trade-offs involved in handling a wider range of prey (Futuyma & Moreno, 1988). However, it is often viewed as an evolutionary dead-end because niche narrowing may limit a species capacity to persist under changing environmental conditions or prey availability, potentially increasing extinction risk (Day et al., 2016). This rationale predicts more evolutionary transitions from generalists to specialists than the reverse. Nevertheless, although specialisation can act as an evolutionary *cul-de-sac* in some instances, most studies find no consistent link with lower diversification rates (Day et al., 2016). Moreover, the specialisation-generalisation continuum is not always clearly delimited. For example, Liem’s Paradox (Liem, 1980) shows that even highly specialised taxa can retain generalist feeding abilities. This apparent contradiction may reflect asymmetric trade-offs, whereby adaptations for capturing difficult or specialised prey do not necessarily constrain the consumption of more common resources. Consequently, specialisation may, in some cases, broaden rather than restrict niche breadth (Robinson & Wilson, 1998).

The study of convergent evolution towards trophic specialisation has often highlighted the predominant role of positive selection acting on protein-coding variants as the main force driving repeated evolution (Roycroft et al., 2021). Furthermore, adaptation is not limited to coding sequences: changes in noncoding regulatory elements can facilitate rapid responses to novel environments (Hao et al., 2023), and gene family expansions or contractions may also contribute to the emergence of adaptive traits (Hoffmann et al., 2010; Roycroft et al., 2021; Whittington et al., 2010). However, selective pressures acting on traits adapted to ancestral niches are expected to be relaxed following the colonisation of new environments, as previously important functions can lose relevance in the new ecological context (Yoder et al., 2010). Moreover, niche specialisation may lead to small, isolated populations with restricted gene flow, potentially decreasing the efficiency of selection (Hung et al., 2017), leading to non-adaptive evolutionary convergence. This raises important questions regarding the genomic signals of convergent evolution: Do convergent lineages exhibit parallel genetic changes fixed by positive selection acting on the same adaptive traits? Or do they instead show signatures of relaxed selective pressures on ancestral traits? Are these changes predominantly affecting proteins or do they involve regulatory modifications? Which is the contribution of gene family evolution to evolutionary convergence? At which biological level does molecular convergence occur? Does it involve changes in the same genes, or does it occur at higher hierarchical levels, such as molecular functions, networks, or wider biological processes? Although genomic data are increasingly available, most of these questions still remain unresolved.

We address some of these questions using the radiation of the spider genus *Dysdera* (Araneae: Dysderidae) in the Canary Islands as a model system, which comprises nearly 60 endemic species derived from a likely single colonisation event (Arnedo et al., 2001, 2007; Bellvert et al., 2023; Macías-Hernández et al., 2016, p. 20212021; Řezáč et al., 2021). Within this radiation, several lineages have independently evolved trophic specialisation (stenophagy) on woodlice, a prey largely avoided by generalist predators due to their morphological, chemical, behavioural defenses, as well as their poorly digestible exoskeleton and high heavy metal content (Drobne, 1997; Schmalfuss et al., 1984; Sutton, 2013; Tuf & Ďurajková, 2022). This specialisation has been accompanied by convergent morphological (Figure 1), behavioural, and physiological changes (Bellvert et al., 2023; Toft & Macías-Hernández, 2017). Despite this biological knowledge, the genomic basis of this dietary specialisation remains largely unexplored.

**Figure 1.**
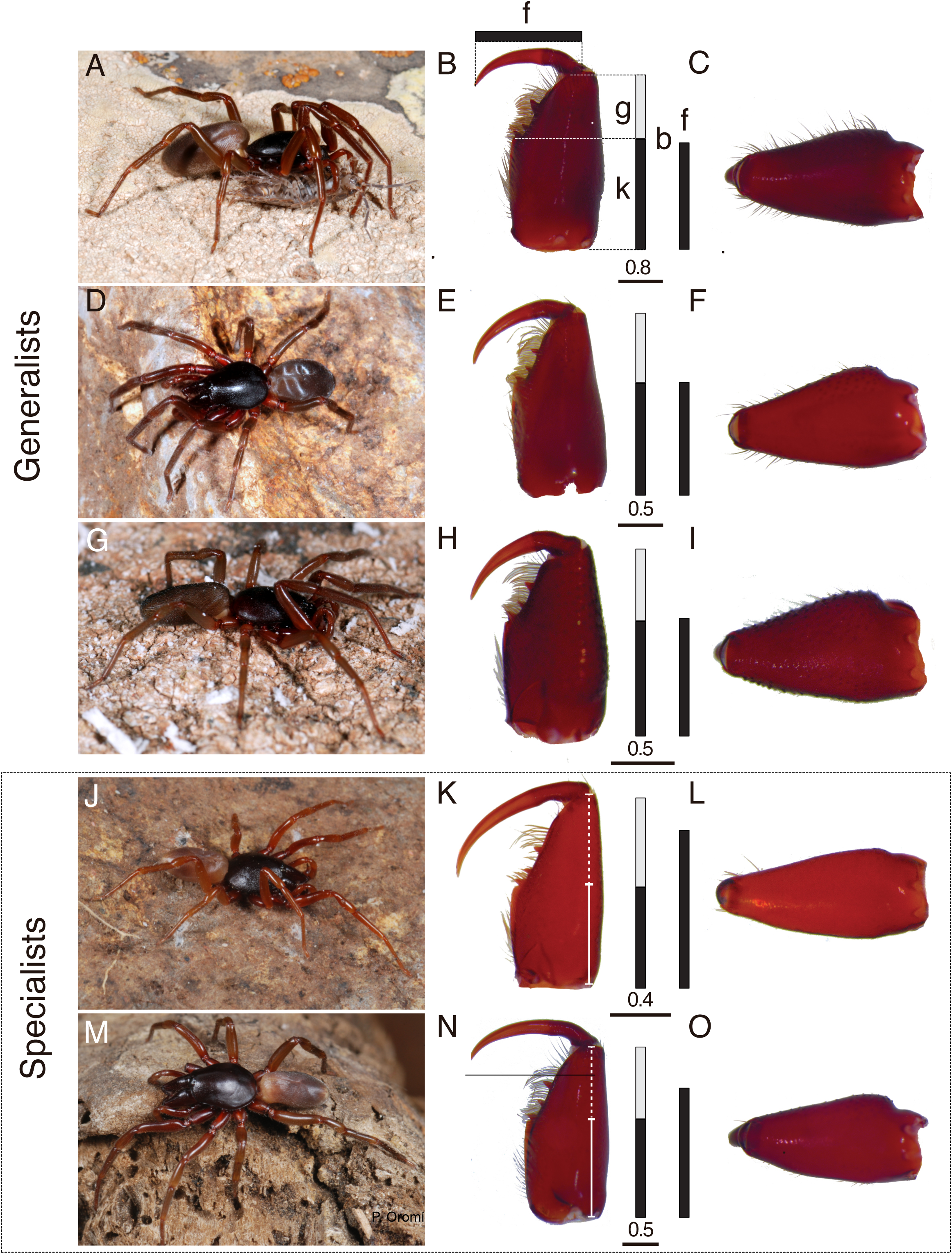
Live images of the target *Dysdera* species (photo credit: Pedro Oromí) and dissecting scope images of the female left chelicera: A–C: *Dysdera silvatica*, La Gomera, A, alive specimen; B, ventral view; C, retrolateral view; D–F: *D. verneaui*, Tenerife, D, alive specimen, E, ventral view, F, retrolateral view; G–I: *D. bandamae*, Gran Canaria, G, alive specimen, H, ventral view, I, retrolateral view; J–L: *D. gomerensis*, La Gomera, J, alive specimen, H, ventral view, I, retrolateral view; M–O: *D. tilosensis*, Gran Canaria, M, alive specimen, N, ventral view, O, retro lateral view. Bars indicate the relative lengths of the different parts of the chelicerae to highlight differences between the standard (generalists) and elongated or slightly elongated (specialists) chelicerae. Left vertical bar: length of the cheliceral groove (g) in white and the internal keel (k) in black, regarding the total length of the basal segment (b). Right vertical bar: length of the cheliceral fang (f). In standard chelicerae, g is approximately 1/3 of b, and f is similar to the length of the internal keel (k), while in elongated chelicerae, g is longer than 2/5 of f, and f is longer than k. Scale bar in mm.

Recent advances in genomics have greatly expanded our ability to investigate the genomic basis of convergent evolution (Vizueta et al., 2019), especially in non-model organisms (Cerca, Cotoras, Santander, et al., 2023; Elmer & Meyer, 2011). These developments enable the identification of genomic changes underlying repeated phenotypic evolution and allow rigorous testing of whether convergence reflects adaptation, relaxation or evolutionary constraints.

To this end, we performed a comprehensive comparative genomic analysis using high-quality genome assemblies of six species of the genus *Dysdera,* three of them newly generated for this study. This dataset enables the comparative analysis of the two independent events of trophic specialisation studied in Vizueta et al., (2019) at the transcriptomic level. As demonstrated by Bellvert et al., (2023), each of these events represents a phylogenetically and geographically independent dietary shift from a generalist ancestor to a specialist descendant lineage (Figure 2). Specifically, we address three questions: (i) whether dietary specialisation is associated with distinct contributions of positive and relaxation of selective pressures, (ii) whether multiple genomic features, including coding sequences, regulatory regions, and gene family dynamics, contribute differentially to this evolutionary transition; and (iii) whether convergent specialisation targets a shared set of genes and functional pathways. By integrating these analyses, we aim to identify the biological processes underlying convergent dietary specialisation and assess its predictability at the genomic level.

**Figure 2.**
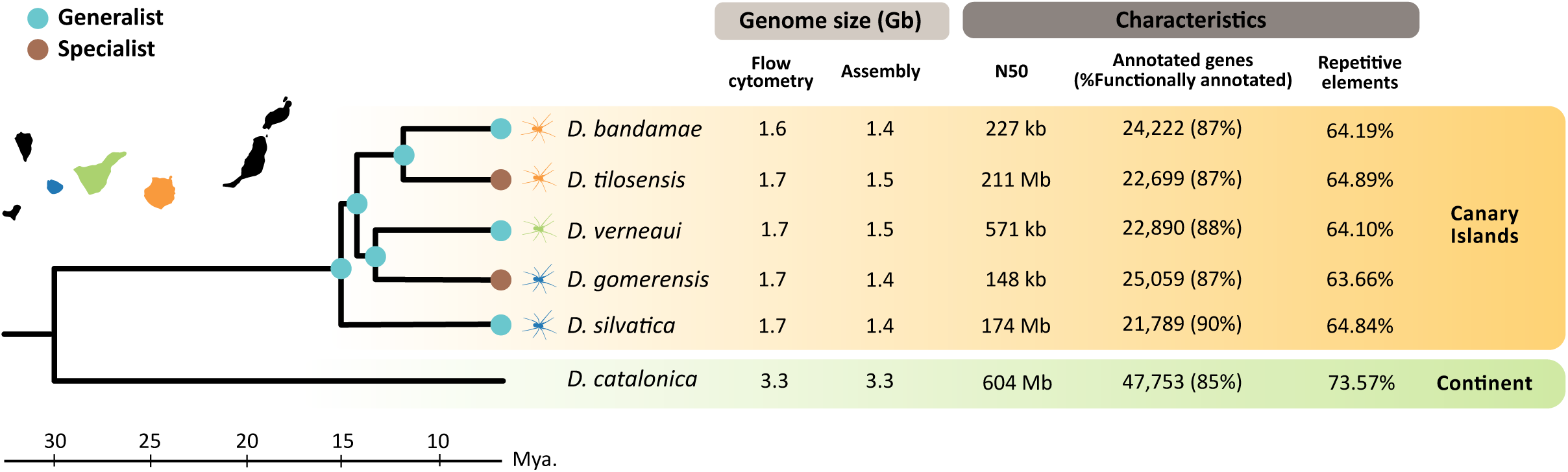
Genome size and assembly statistics for the six surveyed *Dysdera* species. The dated phylogenetic tree was adapted from Crespo et al., (2021) and the references therein. The topology of the Canary Islands endemic clade was independently corroborated by the whole-genome phylogenomic analyses conducted in this study. Coloured circles at the internal nodes and at the tips of the island-endemic species indicate the most probable dietary specialization state inferred from the stochastic character mapping analysis in Bellvert et al., (2023). For the Canary Island species, spider silhouettes indicate the island of origin of the sampled individuals used for genome assemblies (orange: Gran Canaria, green: Tenerife, blue: La Gomera).

## MATERIALS AND METHODS

### Study design, sampling, DNA extraction, sequencing and genome assembly

In this study, we used the genome assemblies of six species of the genus *Dysdera*, five endemics to the Canary Islands (the specialists *D. gomerensis* Strand, 1911, and *D. tilosensis* Wunderlich, 1992, and the generalists *D. verneaui* Simon, 1883, *D. bandamae* Schmidt, 1973 and *D. silvatica* Schmidt, 1981), and one from the European mainland (*D. catalonica* R□ezác□, 2018). Assemblies for *D. silvatica* (Escuer et al., 2022), *D. Tilosensis* and *D. catalonica* Řezáč, 2018 (Pisarenco et al., 2025) were obtained from public databases, whereas those for *D. verneaui, D. gomerensis and D. bandamae* were newly generated in this study (Figure 2). The specimens of these three last species were sampled on the Canary Islands of Tenerife, La Gomera and Gran Canaria, respectively (Supplementary Table 1). The genome sizes of these species were estimated using flow cytometry as described in (Sánchez-Herrero et al., 2019).

DNA extraction was performed using the Gentra Puregene Cell kit (Qiagen). For the PacBio sequencing of *D. verneaui* and *D. gomerensis,* genomic DNA (gDNA) was extracted from a single individual (a male). The small body size of *D. bandamae* precluded the use of a single specimen; therefore, gDNA was extracted from four individuals (two males and two females) to obtain enough gDNA (Supplementary Table 2). The DNA was sequenced using the PacBio CLR technology at Novogene (Cambridge, UK). For the Illumina sequencing, we used DNA from a single individual per species, and all samples were sequenced at Novogene (Cambridge, UK) (Supplementary Table 2).

PacBio reads were used to generate the genome assemblies with wtdbg2 v2.5 (Ruan & Li, 2020), using the realignment mode (–R). PacBio and Illumina reads were aligned to their respective assemblies using minimap v.2.22 (Li 2018) and bwa-mem v0.7.17 (Li & Durbin, 2009), respectively. Polishing was performed in two rounds using the –wtpoa-cns wtdbg2 command, first with long reads and subsequently with short reads. The completeness of the assemblies was determined using the BUSCO-v5.4.2 pipeline (Seppey et al., 2019), with the arachnida (odb10; 2,934 genes), arthropoda (odb10; 1,013 genes), and eukaryota (odb10; 255 genes) datasets (Kriventseva et al., 2019).

To discard potential contaminant sequences, we used the BlobTools2 from BlobTools kit pipeline (Challis et al., 2020; Kumar et al., 2013) with the NCBI nucleotide and UniProt databases (updated in October 2020 and June 2021, respectively) (https://www.ncbi.nlm.nih.gov/nucleotide/, (The UniProt Consortium 2017) UniProt Consortium), along with the previously mentioned BUSCO odb10 databases. Discarded contigs accounted for less than 0.9 % of the total assembly length in all species. Additionally, mitochondrial genome sequences were removed using BLAST+ blastn v.2.12.0 (Altschul, 1997; Camacho et al., 2009) searches against the mitogenome of *D. silvatica* (Adrián-Serrano et al., 2021).

### Repetitive elements, structural and functional genome annotation

Repetitive elements (REs) were masked using RepeatMasker-v4.1.2 (Smit et al., 2013) with the sensitive search (–s), configured with Dfam 3.5 (Storer et al., 2021) and RepBase-20181026 (Bao et al., 2015), and using as input the same RE library as in (Pisarenco et al., 2025). Gene model annotation was carried out with BRAKER-v2.6.4 in ETP mode, integrating protein and RNAseq evidence (Brůna et al., 2021; Hoff et al., 2019; Stanke et al., 2006, 2008). For protein evidence we used the same database as in (Pisarenco et al., 2025). RNAseq reads of each species from (Vizueta et al., 2019) were trimmed with Trimmomatic-v0.39 (Bolger et al., 2014) using the parameters `LEADING:3, TRAILING:3, SLIDINGWINDOW:4:15, MINLEN:50`, and mapped to their respective reference genomes with STAR-v2.7.10a (Dobin et al., 2013). Functional annotation of gene models, was conducted using BLASTP v.2.12 (E-value = 10^-3^) against Swiss-Prot (downloaded in August 2022), and also against a custom arthropoda database (Escuer et al., 2022; Vizueta et al., 2019). Functional annotations were further refined applying InterProscan-v5.57-90.0 with default parameters (Jones et al., 2014) and EggNOG-mapper-v2.1.9 in ultra-sensitive mode (Cantalapiedra et al., 2021; Huerta-Cepas et al., 2017).

### Orthology inference and single-copy orthogroups datasets

Orthogroups across all six species were inferred using Broccoli-v1.2 (Derelle et al., 2020) and OrthoFinder-v2.5.4 (Emms & Kelly, 2019), applying two consecutive rounds of refinement. Briefly, Broccoli was initially used to define the primary orthogroups. Then, genes unclustered by Broccoli were subsequently grouped into additional orthogroups using OrthoFinder.

Single-copy orthogroups generated by Broccoli comprise 4,711 genes in the 5-species Islands dataset (5sp), and 3,567 genes in the 6-species dataset (6sp) which also includes *D. catalonica* used for phylogenetic inference. Coding sequences (CDS) for each gene were then aligned using the codon-wise alignment tool TranslatorX (Abascal et al., 2010). Then, we calculated pairwise genetic distances (divergence) for each gene using Newick Utils v1.6 (Junier & Zdobnov, 2010). Multiple sequence alignments (MSA) including genes with extreme divergence were filtered out, as they likely represent erroneous single-copy inferences or assembling errors. We used the following filtering criteria for the 5-species dataset: (i) a maximum pairwise divergence between any two Canarian species of 0.5, and (ii) a maximum dispersion of divergence values (max - min) of any ingroup species relative to outgroup (*D. silvatica*) lower than 0.25. Per the 6-species dataset, an additional criterion was applied (iii) maximum divergence between the outgroup (*D. catalonica*) and any other species of 0.7. After filtering, the final datasets included 4,463 genes for the 5-species dataset (5sp dataset), and 3,424 for the 6-species dataset (6sp dataset) (Supplementary Table 3). These curated gene sets were used in the downstream analyses.

### Phylogenetic inference

Maximum-likelihood (ML) phylogenies were estimated using IQ-TREE v2.3.6 (Minh et al., 2020) with 10,000 replicates of UFboot and SH-aLRT to assess node supports, and estimating the best-fit evolutionary model through the ModelFinder Plus option (–m MFP). We performed two approaches for both the 5sp and the 6sp datasets: (i) concatenating all MSAs using AMAS (Borowiec, 2016), treating each gene as separate partition, and (ii) inferring a phylogenetic tree for each individual gene, followed by wASTRAL v3 (Zhang & Mirarab, 2022), which accounts for gene tree uncertainty, to infer a species tree from the individual gene trees. Additionally, we quantified the number of gene trees in which the two specialists were recovered as sister lineages using ETE3 Python package (Huerta-Cepas et al., 2016).These complementary approaches allowed us to assess whether the inferred trees were consistent with previously reported phylogenetic relationships among the studied species.

### Selective constraint analyses

We used the HyPhy-v2.5.7 package (Pond et al., 2005) to test for changes in selective constraints in the single-copy orthologs of the 5sp dataset. Specifically, we used BUSTED-PH (Selberg et al., 2026) to identify genes evolving under positive selection associated with dietary specialisation, designating the branches leading to the specialist species (*D. gomerensis* and *D. tilosensis)* as foreground lineages. A gene was classified as a candidate for parallel positive selection when the foreground test (evidence of episodic positive selection in at least one site and one foreground branch) and the foreground–background contrast (selective regime differs between foreground and background branches) were significant after FDR correction (*q-*values < 0.05), whereas the background test (episodic positive selection also occurs on background branches) was non-significant (Selberg et al., 2026). We also used RELAX (Wertheim et al., 2015) to assess potential shifts in selective pressures in foreground lineages, by performing three different background-foreground contrasts (Supplementary Figure 1): (i) with *D. gomerensis* as the foreground branch (Model_g) (ii) with *D. tilosensis* as the foreground branch (Model_t), and (iii) with the lineages leading to both specialist species as foreground branches (Model_sp). We selected genes as candidates for relaxation or intensification when at least one of these three tests was significant (*q-*values < 0.05). For orthogroups with significant *q*-values in more than one comparison, the model with the lowest Akaike Information Criterion (AIC) was selected as the best fit model. Candidates were further classified as relaxed if *k* < 1, or under intensification if *k* > 1. Although selection intensification inferred by RELAX has a more complex interpretation than a straightforward signal of positive selection, we conservatively considered as candidates for parallel positive selection only those orthogroups supported by both RELAX (*k* > 1) and BUSTED-PH (see above). For these orthogroups, we considered the best-fitting model inferred by RELAX. To assess the chromosomal location of genes under distinct selection regimes, we used *D. tilosensis* annotations as the reference.

### Evolutionary rate shift analysis in the 5’ upstream gene regions

For this analysis, we focused on 1 kb regions upstream of single copy-orthologs in the 5sp dataset, as this length has commonly been used in previous studies of arthropods and nematodes (Chesters et al., 2026; Liu et al., 2004). First, we extracted the 1 kb upstream gene regions using custom Python scripts incorporating BEDTools-v2.30 functions (Quinlan & Hall, 2010). Specifically, we used flank –s to define strand-specific upstream regions, subtract to discard upstream regions overlapping adjacent annotated genes, and getfasta –s to retrieve strand-oriented upstream regions in FASTA format. Only orthogroups with full-length upstream regions for the five species were included in further analyses (u5sp_1 dataset).

The upstream regions of each gene were aligned using the –linsi algorithm of MAFFT-v7.310 (Katoh & Standley, 2013) to ensure high-accuracy alignments despite potential high sequence divergence. To detect parallel evolutionary rate shifts in specialist lineages, we employed PhyloAcc-GT (Yan et al., 2023), which accounts for loci tree discordance. We used the –filter option to exclude MSA containing excessive gap content. Default logBF cutoffs (i.e., logBF1 > 1 for evidence of acceleration in foreground lineages, and LogBF2 > 1 for evidence that this acceleration is specific of these lineages), were applied to select the best fit model for each orthogroup. The trees based on 5 species from wASTRAL were used as input for PhyloAcc-GT. This software used neutral background tree and the corresponding transition rate matrix inferred from the 4-fold degenerate sites in the concatenated alignment of the orthogroups from the 5sp dataset, build with phyloFit program from the PHAST v1.4 package (Hubisz et al., 2011).

### Estimation of gene family turnover rates

We used BadiRate v.1.35 (Librado et al., 2012) to obtain maximum likelihood (ML) estimates of gene family turnover rates under the birth-death-innovation model (BDI). We compared five different branch-specific models: (i) the global rates model (GR), where all branches evolve under the same turnover rates; (ii) the free rate model (FR), which assumes independent rates for all branches; (iii) the specialist-specific model (spBR), that assumes that the two specialist lineages share gene family turnover rates that differ from those of the background lineages; (iv) *D. gomerensis*-specific rate model (gBR), assuming that *D. gomerensis* has turnover rates that differ from the rest of lineages and, (v) the equivalent species-specific model but designating *D. tilosensis* as the focal lineage. For these analyses, we used the OG_5sp dataset and an ultrametric species tree obtained applying the chronos function from the R package ape (Popescu et al., 2012) to the phylogenetic tree inferred from the 5sp dataset. As the spBR model showed the best support (Supplementary Table 4), we therefore performed further analyses using the spBR model. After identifying the best-fitting model of genome-wide gene turnover rates, we analysed separately each orthogroup (using the OG_5sp_2 dataset) to identify those families with accelerated rates in specialist lineages. To infer gene family expansions and contractions in specialists and generalists we analysed each orthogroup under the spBR model defining three sets of test branches: terminal specialist branches, terminal generalist branches, and a background set comprising all remaining ones.

### GO enrichment analyses

To assess whether candidate genes of the different comparative genome analyses were overrepresented in specific GO categories, we performed a series of GO enrichment tests. Firstly, we obtained the GO terms for annotated genes of the five species using InterProScan v.5.57–90.0 (Jones et al., 2014). Functional annotations were mapped at the orthogroup level by pooling the GO terms of all orthologous genes. Within each orthogroup, redundant terms were discarded to provide a non-redundant dataset for the GO enrichment analysis. For tests involving the 5sp dataset (positive selection and relaxed selection), all orthogroups identified in this dataset, together with their associated GO terms, were used to define the reference universe. For the GO enrichment of the upstream region rate shifts analysis, this reference universe was restricted to orthogroups for which the full upstream length was successfully retrieved across all species. Moreover, we evaluated three sets of candidate genes (Supplementary Table 5) (i) those identified in both specialist lineages, (ii) those only found in *D. gomerensis* or (iii) only in *D. tilosensis*. For the GO enrichment analyses associated with gene family turnover rates, the universe set comprised all orthogroups inferred with Broccoli and Orthofinder. In this case, we tested two candidate gene sets: the top 10% orthogroups with the highest birth rates and the top 10% with the highest death rates.

The GO enrichment analysis was performed using the R packages GSEABase v.1.60.0 and GOstats v.2.64.0 (Falcon & Gentleman, 2007; Morgan et al., 2022). We ran GOstats applying the conditional testing option and considered GO terms significant when *p*-value < 0.01 and were present in at least five genes. We also used the R packages clusterProfiler v.4.8.1 (Wu et al., 2021) and AnnotationForge v.1.42.0 (Carlson & Pagès, 2024) to transform the GOstats results into an enrichResult object, and visualised the results using the R packages enrichplot v.1.20.0 (Yu, 2023) and ggplot2 v.3.4.2 (Wickham, 2016).

## RESULTS

### New genomic resources confirm previously reported phylogenetic relationships

We first established a comprehensive genomic and phylogenetic framework for studying focal species. We generated new genome assemblies of three Canary Islands endemic species of the genus *Dysdera*: *D. verneaui, D. bandamae*, and *D. gomerensis* (Supplementary Table 1). Flow cytometry estimates showed haploid genome sizes of approximately 1.7 Gb for all three species (Figure 2, Supplementary Table 2), consistent with previously reported values for the two other Canary Island species, *D. silvatica* and *D. tilosensis* (Pisarenco et al. 2025). Estimates from genome assemblies matched those obtained from flow cytometry estimates (Figure 2, Supplementary Table 2). Genome completeness is high in all species, with >95% of Eukaryota BUSCOS recovered, and >85% complete BUSCOs in the Arachnida, Arthropoda and Eukaryota datasets (Figure 2, Supplementary Table 6, Supplementary Table 7). Regarding the three new assemblies, *D. verneaui* (N50 = 571 kb) is the most continuous, followed by *D. bandamae* (227 kb) and *D. gomerensis* (148 kb); these values are relatively high for non-chromosome-level assemblies (Supplementary Table 6). In line with previous estimates for *D. silvatica* and *D. tilosensis*, repetitive elements (REs) account for a large proportion of their genome sequences (around 64% in all three species (Figure 2, Supplementary Table 6). Structural annotations identified between 22,890 and 25,059 protein-coding genes (Figure 2, Supplementary Table 6), of which around 87% were functionally annotated.

Phylogenetic analyses using new genomic sequences corroborate the relationships reported in previous transcriptomic (Vizueta et al., 2019) and mitogenomic (Adrián-Serrano et al., 2021) studies (Supplementary Figure 2; Supplementary Figure 3), recovering a clade comprising two generalist-specialist sister-species pairs (*D. bandamae - D. tilosensis,* and *D. verneaui - D. gomerensis*) with *D. silvatica* as sister taxon of the remaining island species. Moreover, specialist species were recovered as sister lineages only in 8.4% and 6.6% of single-gene phylogenetic trees from the 5sp and 6sp datasets, respectively, with low bootstrap support for this node (mean bootstrap of 54.3 and 53.9, respectively). This result supports the independent phylogenetic origin of focal specialist lineages, and that incomplete lineage sorting (ILS) is unlikely to play a major role in the evolution of these two lineages. Crucially, mapping the ancestral-state reconstructions of cheliceral morphotypes inferred by stochastic character mapping (Bellvert et al., 2023), which directly predict specialist and generalist feeding strategies, onto our phylogeny provides strong support for two independent origins of dietary specialisation.

### Specialists show convergent signals of positive selection

Among the 4,463 orthogroups analysed with RELAX, 523 showed significant signals of parallel relaxation in specialists (model_sp; Figure 3a, Table 1), and 208 and 183, are relaxed exclusively in *D. gomerensis* and *D. tilosensis*, respectively (Figure 3a, Table 1, Supplementary Table 8). The ratio of intensified-to relaxed-selection candidates is high in those comparisons where species-specific models are the best-fitting models (1.68 in Model_g and 1.89 in Model_t). This ratio is considerably lower in comparisons were the data fit better the Model_sp (0.58), suggesting that relaxation predominates when changes in selective constraint occur in parallel across the two specialist lineages (Table 1, Figure 3a, Figure 3b). We then used BUSTED-PH to identify genes under positive selection associated with the specialist diet. We found that most genes showing signatures of positive selection in specialists overlapped with those identified in RELAX as undergoing intensification (from 97.7 to 99 %). Among them, 102 showed parallel signatures in specialist species, and 86 and 88, were exclusive to *D. gomerensis* or *D. tilosensis*, respectively (Figure 3c, Table 1, Supplementary Table 9). Genes under intensified and relaxed selection were relatively evenly distributed across chromosomes (Supplementary Figure 4).

**Figure 3.**
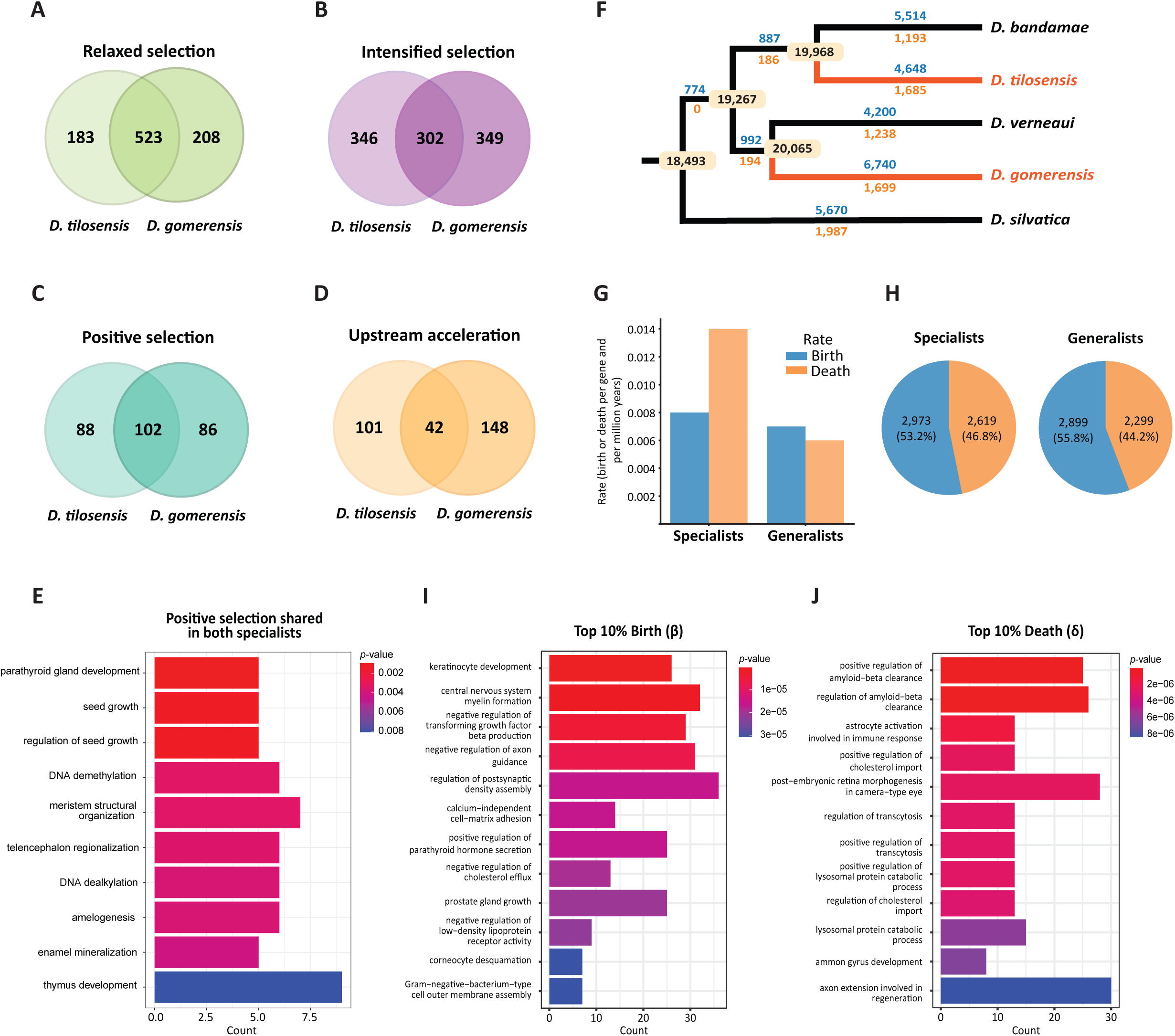
A, Number of genes under relaxed selection; B, intensified selection; C, positive selection. D, Number of upstream accelerated regions. E, GO enrichment analysis of significantly enriched BP (biological processes) terms (*p*-value < 0.01) for positively selected genes shared by both specialists (Model_sp). F, Estimation of gene family turnover rates using the Badirate BDI model, allowing distinct rates for specialists based on 29,586 orthogroups. The phylogenetic tree shows the gene gains (blue) and losses (orange) along each branch, with ancestral gene counts indicated in beige boxes. Branches of specialist lineages are highlighted in orange. G, Estimated birth (blue) and death (light orange) rates per gene and per million years for each branch category. H, Comparison of the number of orthogroups in which either birth (blue) or death (light orange) is the highest rate in specialists versus generalists lineages. I, GO enrichment analysis of significantly enriched BP terms (*p* < 0.01) for the top 10% orthogroups with the highest estimated birth rates (I), or the top 10% death rates (J) in specialist lineages.

**Table 1.**
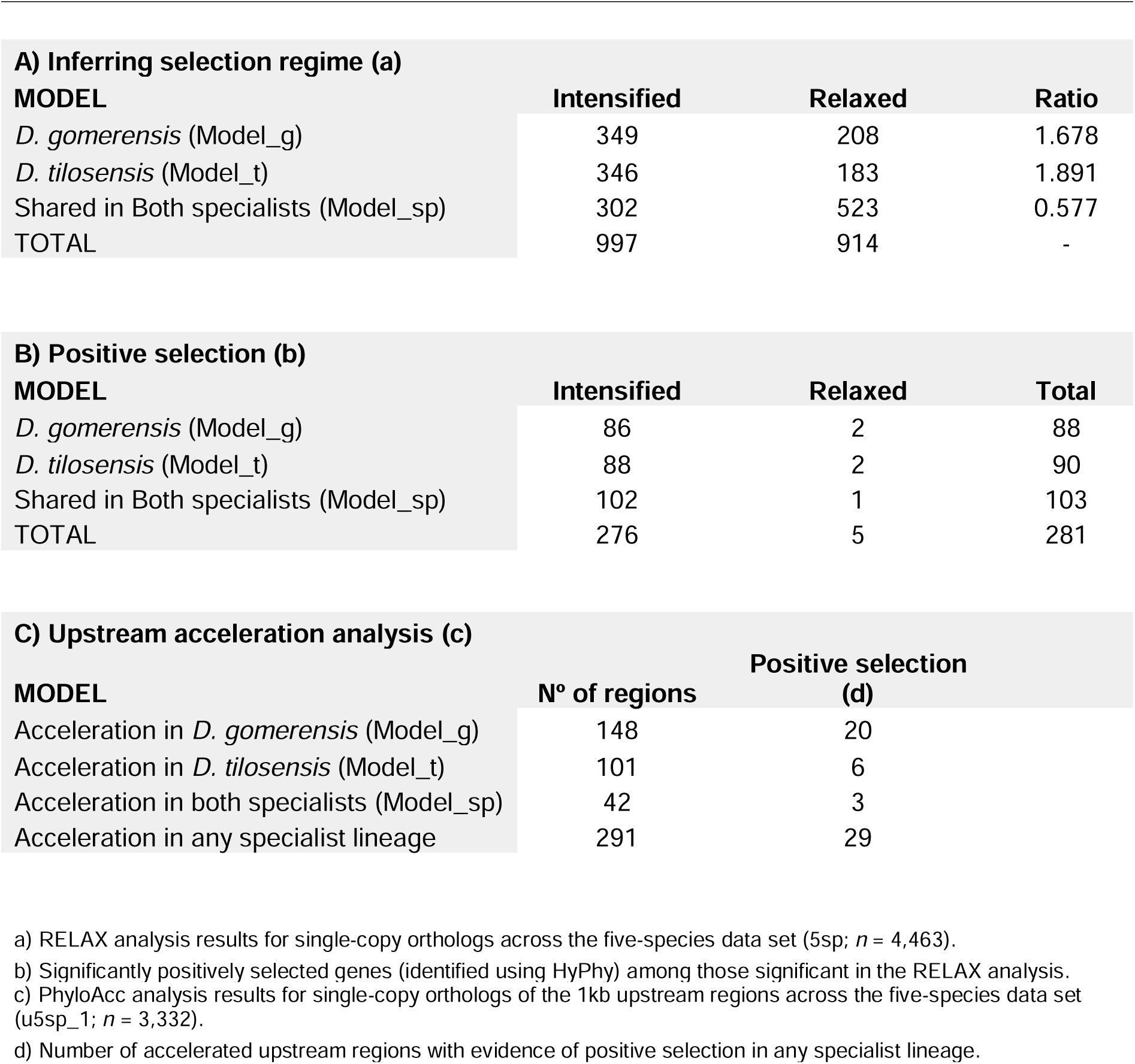
Summary of significant genes and regions under different selective constraints.

The biological processes (BP) significantly enriched (*p* < 0.01) among genes under positive selection in specialists were related to Gene Ontology (GO) terms associated to stress responses and DNA methylation (Figure 3e, Supplementary Table 10). Particularly, the genes showing parallel signals of positive selection in specialists are enriched in GO terms associated to gene expression, such as DNA demethylation and dealkylation, whereas specialist specific selection targets are related to histone H4 acetylation and protein autoubiquitination (a stress response involved in damaged protein degradation) in *D. tilosensis*, and stress response, proteoglycan and glycoprotein metabolic and biosynthetic processes in *D. gomerensis*. In this last species, molecular functions (MF) related to transferring sulphur-containing groups and iron ion binding are also enriched among positively selected genes (Supplementary Figure 5).

We also found stress response to copper ions, cytokine immune response and tumour necrosis factor signalling to be significantly enriched in the genes showing parallel relaxation in both specialists (Supplementary Figure 6, Supplementary Table 11), although these last two terms are annotated primarily from vertebrate data and should therefore be interpreted cautiously. When considering each specialist species separately, *D. tilosensis*, relaxed genes were mostly enriched in GO terms associated with meiotic cytokinesis, microtubule polymerization and cytoskeleton-dependent cytokinesis, as well as cellular component (CC) terms related to phagocytic and Golgi-associated vesicles. In *D. gomerensis*, significant relaxed genes are enriched in adult chitin-based cuticle development and neurologically associated CC terms such as glial cell projection.

To identify potential regulatory changes associated with dietary specialisation, we screened 3,332 upstream gene regions (1 kb) for signatures of accelerated evolution. Of these, 42 showed acceleration in the two specialists, 148 exclusively in *D. gomerensis*, and 101 exclusively in *D. tilosensis* (Figure 3d, Table 1, Supplementary Table 12). Among the total of regions with accelerated upstream regions, 29 were also identified as being under positive selection. As observed for intensified and relaxed genes, accelerated regions are broadly distributed across chromosomes (Supplementary Figure 4). The GO enrichment analysis of the 42 genes associated with upstream regions accelerated in both specialists did not recover any significantly enriched GO terms (Supplementary Table 13). In contrast, genes with upstream regions accelerated exclusively in *D. gomerensis* (Model_g) were enriched in BP-related to oxidative stress, such as response to genistein, aminoglycan metabolic process and response to hydroxyisoflavone. In terms of MF, enrichment was observed for chitin binding (Supplementary Figure 7, Supplementary Table 13). In *D. tilosensis* (Model_t), enriched functions were mostly neurological, such as regulation of smooth muscle cell migration and establishment of the blood-brain barrier (Supplementary Figure 7, Supplementary Table 13). Additional enrichment was observed for processes related to larval salivary gland morphogenesis. Consequently, we found limited signal of convergent regulatory evolution either at the level of individual genes or at broader hierarchical levels.

### Gene families exhibit increased death rates in specialist lineages

To explore the gene gain and loss dynamics associated with dietary specialisation, we analysed gene family turnover rates in the focal genomes. Overall estimations of gene turnover rates indicate that data are better fitted by the specialist-specific model (BRsp) (Supplementary Table 4). Remarkably, while generalist and specialist species exhibit similar gene birth rates (0.007 and 0.008 gains per gene and per million years, respectively), the gene death rate in specialists is more than twice that of generalists (0.006 in generalists vs. 0.014 in specialists), see Figure 3f, Figure 3g, Figure 3h, Supplementary Table 4).

We determined the functional profile of genes undergoing rapid expansion or contraction associated with diet specialisation analysing the 10% orthogroups with the highest birth and death rates in specialist lineages (Supplementary Table 4, Supplementary Table 14, Supplementary Table 15; Figure 3i, Figure 3j, Supplementary Table 16). We found that orthogroups with the highest birth rates were mainly related to the nervous system, including the negative regulation of transforming growth factor beta production (which influences amyloid beta formation), and the negative regulation of axon guidance. Other enriched BP are related to cholesterol efflux, and calcium associated processes, such as parathyroid hormone secretion (which regulates calcium levels) and calcium independent cell-matrix adhesion. We also detected epidermal processes, including keratinocyte development and corneocyte desquamation, and mercury ion binding as enriched MF. However, these GO terms should again be interpreted with caution because they are vertebrate specific. Orthogroups with the highest death rates were enriched in BP related to positive regulation of amyloid-beta clearance and cholesterol import. We also detected additional enriched GO terms related to vesicular transport across the cell and extracellular cargo degradation, including trancytosis and lysosomal-associated processes, as well as axon regeneration functions. Our results show that rapid gene family turnover in specialists affects different physiological processes: expansions are enriched for nervous system and cholesterol regulation families, whereas contractions primarily impact vesicular transport and nervous system genes.

## DISCUSSION

### Parallel genomic changes were dominated by relaxed selection and gene family contractions

Convergent evolution is often associated with adaptation, although similar phenotypes may arise through distinct evolutionary forces. While adaptation to new ecological niches is typically linked to positive selection acting on protein-coding and regulatory regions (Biswas & Akey, 2006; Hao et al., 2023; Roycroft et al., 2021), traits that lose functional relevance in novel environments may experience relaxed selection, leading to functional decay or gene loss (Yoder et al., 2010). In this context, gene family dynamics provide additional insight into convergent evolution (Hoffmann et al., 2010; Roycroft et al., 2021; Whittington et al., 2010). Indeed, redundancy among paralogous gene copies make gene families particularly suitable for detecting signatures of selection. Nevertheless, dietary specialisation is commonly expected to lead to gene family contraction, as gene copies required for exploiting a broader diet may become dispensable after specialisation (Hecker et al., 2019). Although gene loss can occasionally be adaptive (Steindorff et al., 2024), it more frequently reflects relaxed selection (Sharma et al., 2018), with potentially long-term evolutionary consequences. Consistent with this framework, we detected a higher proportion of relaxed genes among the parallel responses in specialists, which align with the higher rates of gene loss also found in these lineages (Table 1, Figure 3a, Figure 3B, Figure 3G). Conversely, lineage-specific changes were more frequently under intensified selection than parallel changes. Although intensification of selection does not necessarily indicate adaptation, a higher proportion of positively selected genes were also identified as under intensified selection (Table 1).

Together, our results show that parallel genomic responses are predominantly characterised by relaxation of selective constraint and gene family contractions likely associated with the loss of functions required under the ancestral generalist diet, whereas stronger signals of positive selection, selection intensification, and gene family expansion are associated with the refinement of specialist functions at different hierarchical levels.

### Key adaptations underlying trophic specialisation

Remarkably, we found that positively selected genes in specialists are primarily associated with methylation processes and stress responses, consistent with heavy metal tolerance mechanisms reported in previous studies (Guan et al., 2019; Janssens et al., 2009), and in our transcriptomic analyses in the same species (Vizueta et al., 2019). These processes are typically triggered by metal-induced oxidative stress, which disrupts cellular homeostasis and require activation of molecular pathways to maintain cellular integrity (Korsloot, 2004). Metal-induced stress also alters DNA methylation patterns (Guan et al., 2019) and, in collembola, similar shifts in transcriptional regulation have been suggested to enhance metal tolerance (Janssens et al., 2009). The signal of positive selection detected on genes involved in histone H4 acetylation in *D. tilosensis* strengthens this idea, as nickel exposure is known to affect this process in humans (Martinez-Zamudio & Ha, 2011). This supports the hypothesis that positive selection enhances detoxification capacity in specialist *Dysdera* to tolerate high concentrations of toxic heavy metals bioaccumulated in woodlice. Additionally, enriched functions linked to protein degradation such as ubiquitination (notably in *D. tilosensis*) may contribute to the excretion of damaged or heavy-metal-carrying proteins (Ursu & Zhitkovich, 2026). On the other hand, *D. gomerensis* may rely on an alternative detoxification mechanism, as genes associated with proteoglycan-related functions were found to be under positive selection, and their glycosaminoglycans (GAGs) components possess metal-chelating properties in synthetic chemistry assays (Shitrit et al., 2022). Given that chelation is a well-established medical strategy for reducing heavy metal toxicity (Flora & Pachauri, 2010), these results raise the possibility that a similar chelating response coupled with the stress-response mechanism, contributes to mitigate metal toxicity in *D. gomerensis*. Together, these results suggest that while both specialists share core adaptive functions related to transcriptional regulation and stress to chemical pressures, species-specific signals suggest divergent evolutionary solutions: *D. tilosensis* shows broader physiological reconfiguration, whereas *D. gomerensis* exhibits more stress responses. However, it remains to be determined whether these alternative adaptive strategies are associated with the differences in levels of stenophagy or in cheliceral shape observed between the two species (Bellvert et al., 2023). Overall, our results suggest that positive selection may promote enhanced heavy metal detoxification, likely by targeting pathways involved in metal excretion or chelation and cellular stress response. These findings are consistent with earlier toxicology studies in *D. crocata*, which show that these spiders retain only a fraction of ingested metals and do not accumulate high heavy metal levels even when fed isopods with varying metal concentrations, suggesting an efficient excretion mechanism (Hopkin & Martin, 1985).

In contrast, relaxed genes are enriched for functions related to the response to the copper ion stress, one of the metals known to bioaccumulate in isopods (Prosi & Dallinger, 1988). Although this finding may initially seem counter-intuitive, it could instead reflect the efficient detoxification mechanisms discussed above. In specialist species, if heavy metal toxicity is efficiently mitigated through enhanced detoxification mechanisms, genes involved in the canonical heavy metal stress response may become less critical, leading to reduced selective constraint and consequently, relaxed selection, potentially reducing the physiological costs of stress-response activation. Additionally, in *D. tilosensis,* the observed reduced investment in cytoskeleton-dependent cytokinesis, involved in cell division, is consistent with previous evidence (Jomova et al., 2025; Lv et al., 2023; Vizueta et al., 2019; Wang et al., 2018).

In upstream regions, no shared functional enrichment is detected in both specialists, suggesting that acceleration in upstream, putative regulatory regions, does not act on common pathways; instead, they are largely lineage-specific. On the one hand, *D. gomerensis* shows acceleration in upstream regions of genes related to oxidative stress, which may act in concert with those detected as positively selected in stress responses identified in this lineage (Janssens et al., 2009; Korsloot, 2004; Mijin et al., 2023; Zhu et al., 2016). On the other hand, *D. tilosensis* is enriched for neurological functions, including blood-brain barrier-related genes that may mitigate the increased permeability neurotoxic effects associated with heavy metal accumulation (Zahoor et al., 2024). Furthermore, acceleration in upstream regions of genes associated with salivary gland-related genes observed in *D. tilosensis* mirrors patterns observed in centipedes exposed to long-term cadmium, where storage-related organelles are reduced in this cell type (Rost-Roszkowska et al., 2020). These changes might represent compensatory adaptations to mitigate similar toxic effects, given that cadmium is commonly bioaccumulated by isopods (Prosi & Dallinger, 1988).

### Mirrored Functional Patterns in Expanded and Contracted Gene Families

Gene families exhibit a significant twofold increase in gene death rates in specialist species supporting a model of genomic streamlining in which functions no longer required for specialised diets are lost. The fact that this occurs in a multigene family context would indicate that some paralogs become dispensable, but others are still needed to preserve limited dietary flexibility. Notably, gene families involved in vesicular transport and lysosomal processes show strong contraction, consistent with previous transcriptomic evidence (Vizueta et al., 2019), and also with functional categories enriched among genes relaxed in *D. tilosensis*. Different organisms employ diverse strategies to cope with heavy metal exposure (Ahearn et al., 2004). Metallothioneins can sequester metals as a short-term tolerance mechanism, after which metals may be excreted or stored in vesicular compartments, such as specific granules and lysosomes (Vijver et al., 2004). This latter long-term tolerance strategy (i.e storage) is common in collembola (Vijver et al., 2004) and crustaceans (Ahearn et al., 2004), while isopods bioaccumulate metals in the hepatopancreatic granules (Prosi & Dallinger, 1988). In contrast, the contraction of gene families involved in vesicular transport and lysosomal processes suggests that specialist *Dysdera* likely rely less on vesicle-mediated metal storage and more on rapid metal excretion. Although the mechanisms of heavy metal detoxification in spiders remain unclear and vary among species (Yang et al., 2016), our interpretation aligns with findings of some spider species regulating the excess of heavy metals in food by efficient release in faeces (Wilczek & Babczyńska, 2000) and with previous observations in *D. crocata* showing no increase in bioaccumulated heavy metals, pointing to an expulsion mechanism (Hopkin & Martin, 1985).

Despite the overall acceleration of gene loss rates in specialists, specific gene family expansions can also point to adaptive responses. The expanded families include functions related to metal-binding and epidermal processes, which may contribute to oxidative stress mitigation (Janssens et al., 2009). For instance, enriched epidermal-related functions could contribute to toxicity mitigation, potentially linked to moulting-mediated metal removal, as suggested in *D. crocata* (Hopkin & Martin, 1985). Similar exoskeleton bioaccumulation-based detoxification strategies have been reported in other organisms, including woodlice (Raessler et al., 2005), crabs (Bergey & Weis, 2007; Butler & Zou, 2021), collembola (Kristiansen et al., 2021) and crayfish (Xiong et al., 2021). However, as *Dysdera* cease moulting after reaching adulthood, as is typical in most spiders (Guarisco, 1991), moulting is unlikely to represent their primary detoxification mechanism in adult individuals. Consistent with this, we found relaxation of selection on genes involved in adult chitin-based cuticle development in *D. gomerensis*, which is inconsistent with a prominent role for moulting in detoxification.

Remarkably, functional enrichment further reveals a mirrored pattern: some gene family expansions are accompanied by contractions of those with overlapping roles. This pattern is particularly evident in amyloid-beta formation and cholesterol metabolism, where expansions of cholesterol efflux-related genes are accompanied by contractions in import-related ones. This suggests a potential overall reduction in cholesterol levels. This finding is particularly relevant given that heavy metals exposure can increase cholesterol levels (Jomova et al., 2025), and that both, heavy metals and cholesterol metabolism, contribute to amyloid-beta plaque formation (Bishop & Robinson, 2004; Mijin et al., 2023; Rudajev & Novotny, 2023). Consistently, we observed gene family expansions associated with negative regulation of amyloid-beta production, which may further limit plaque formation. Following this rationale, reduced amyloid-beta levels may decrease selective pressure for clearance mechanisms, consistent with the observed contraction of gene families involved in this function. Additionally, as heavy metal exposure can reduce axon length (Zhu et al., 2016), the expansion of genes involved in the negative regulation of axon guidance may reflect a mechanism to halt growth in toxic conditions. Under chronic heavy metal stress, contraction of gene families associated with axon regeneration could be adaptive, as reducing ineffective repair processes may minimise metabolic costs.

### Proposed Mechanisms for the Adaptation of Specialist *Dysdera* towards an Isopod-Rich Diet

Taken together, our results lead us to propose the hypothesis that specialist *Dysdera* have evolved targeted genomic adaptations enabling efficient detoxification and survival on heavy metal-rich diets. These adaptations appear to operate at multiple levels: (i) mitigating heavy metal-induced neurotoxicity, (ii) reducing reliance on intracellular storage of toxic compounds, (iii) enhancing efficient metal excretion, potentially through some yet-uncharacterised expulsion mechanism. Some of these adaptations appear to be driven by positive selection acting on protein-coding genes involved in stress responses, likely enhancing detoxification capacity. This may be further reinforced by our results showing that gene families related to mercury ion binding would also have been targets of adaptive expansions. Indeed, gene family dynamics further suggest complementary mechanisms of neural protection, including processes that directly reduce amyloid-beta plaque formation and indirectly regulate cholesterol levels as putative targets of selection. These processes, which are conserved between arachnids and humans, are particularly relevant under metal-induced stress and widely studied in the context of Alzheimer’s disease (Fabian-Fine et al., 2024; Qi et al., 2023). Moreover, lipid gene family changes in specialist *Dysdera* may reflect the need for its efficient processing, given that lipids are primarily obtained from generalist prey rather than from isopods (Toft & Macías-Hernández, 2021). Consistent with a detoxification strategy based on rapid excretion, contractions in gene families involved in lysosomal, vacuolar and transcytosis pathways suggest reduced investment in intracellular storage and vesicle-mediated transport, which is supported by toxicological evidence in *D. crocata* (Hopkin & Martin, 1985). Accelerated evolution in upstream regulatory regions suggests lineage-specific optimisation, with *D. gomerensis* showing signatures linked to oxidative stress responses, and *D. tilosensis* to neurological protection, including reduced blood-brain barrier-related functions. However, as those signals are not detected in both specialists, it is uncertain that these functions are truly related to dietary specialisation.

The differences in the genomic signals of specialisation found in *D. gomerensis* and *D. tilosensis* might reflect variation in island-specific ecological conditions and/or demographic histories. For instance, Gran Canaria is larger and potentially more ecologically diverse than La Gomera. The slightly higher ratio of intensified to relaxed selection in *D. tilosensis* may indicate greater selection efficiency, potentially associated with a larger effective population size, whereas the stronger signature of relaxation in *D. gomerensis* may reflect a greater influence of genetic drift. However, these interpretations remain speculative and require explicit demographic inference.

Several limitations of this study should be acknowledged. First, functional inferences based on GO annotations are constrained by using a non-model organism, as many genes lack GO annotations, and available annotations are largely transferred from model species. Accordingly, our functional interpretations should be regarded with caution. Experimental validation will therefore be essential both to corroborate our findings and to refine functional annotations. In addition, extending these analyses to a broader range of independently evolved specialist lineages would provide a more robust assessment of the generality and repeatability of the genomic patterns reported here.

Overall, our study uncovers the genomic signatures of the convergent dietary specialisation. Positive selection targets a small core set of functionally convergent adaptive processes, primarily related to heavy metal tolerance, at different biological hierarchical levels. In contrast, the relaxation of selective constraints affects genes that have likely become dispensable under the new dietary regime. Together, these findings suggest that similar ecological pressures would have driven convergent, yet non-identical, molecular solutions, supporting the idea that dietary specialisation follows partially predictable evolutionary trajectories.

## Author Contributions

J.R., S.G.-R., and M.A.A. conceived the study. A.S.-G. contributed to the design of the selected analyses. M.O.-M., V.A.P. and C.F.-L. performed the analyses. N.M.-H. and M.A.A. participated in the sampling campaign. M.O.-M., J.R., S.G.-R. and A.S.-G. interpreted the data. M.O.-M. drafted the first version of the manuscript. All authors revised and approved the final manuscript.

## Acknowledgements

This work was supported by Ministerio de Ciencia e Innovación (MCIN/AEI/10.13039/501100011033) grants (PID2019-103947GB, PID2019-105794GB, PID2022-137758NB-I00, PID2022-138477NB, and an FPI fellowship to V.A.P. PRE2020-095592). Additional funding was provided through a predoctoral fellowship from the Universitat de Barcelona to M.O-M. (PREDOCS-UB 2021), and by Comissió Interdepartamental de Recerca i Innovació Tecnològica (2021SGR00279, 2021SGR0689). We gratefully acknowledge the Cabildo of Gran Canaria and the Canarian Government for granting collection permits and providing support with accommodation and logistics during fieldwork. We thank Carla Díaz, Salvador de la Cruz, Luís Crespo and Adrià Bellvert, who assisted in the field work. We thank Daniel Dols-Serrate for his valuable comments and his help in the phylogenetic inference analyses.

## Data Availability

The three whole-genome nucleotide sequences have been deposited at DDBJ/ENA/GenBank under Bioproject accession number PRJNA1463000, and BioSample accession numbers SAMN59219342-SAMN59219347 and SAMN59219375-SAMN59219377. Structural and functional annotations are available in github: https://github.com/molevol-ub/Dysdera_BanGomVer_genomes

## Ethical statement

The study involved the collection of spiders from natural populations for molecular analyses. All specimens were collected in accordance with applicable national and local regulations, and all required governmental permits were obtained prior to sampling. No experiments involving the manipulation or treatment of live animals were conducted, and therefore no specific ethical approval for animal experimentation was required. The study complied with all applicable legislation and guidelines governing the collection and use of specimens from natural populations.

## Supplementary Tables

Supplementary Table 1. New samples used in this study

Supplementary Table 2. New DNA-RNA sequence data

Supplementary Table 3. Summary of the orthogroup datasets

Supplementary Table 4. Gene family turnover rates using BadiRate

Supplementary Table 5. Summary of the GO enrichment input datasets

Supplementary Table 6. Genome assembly and annotation statistics

Supplementary Table 7. BUSCO results

Supplementary Table 8. Relaxed and intensified orthogroups (RELAX results)

Supplementary Table 9. Positively selected orthogroups (BUSTED-PH results)

Supplementary Table 10. GO enrichment analysis of positively selected genes

Supplementary Table 11. GO enrichment analysis of genes under relaxed selection

Supplementary Table 12. Accelerated upstream regions analysis using PhyloAcc-GT

Supplementary Table 13. GO enrichment analyses of the upstream accelerated regions

Supplementary Table 14. Top 10% orthogroups with highest birth rates

Supplementary Table 15. Top 10% orthogroups with highest death rates

Supplementary Table 16. GO enrichment analysis of the expanded and contracted gene families

## Supplementary Figures

Supplementary Fig. 1. Tested models in the selective constraint analysis.

Supplementary Fig. 2. Phylogenetic inference of the 5 species of the Canary islands.

Supplementary Fig. 3. Phylogenetic inference of the 5 species of the Canary islands and the mainland species *D. catalonica*.

Supplementary Fig. 4. Chromosomal distribution of genes under the different selective regimes analysed.

Supplementary Fig. 5. GO enrichment analysis of the positively selected genes exclusive of *D. gomerensis* and exclusive of *D. tilosensis*.

Supplementary Fig. 6. GO enrichment analysis of genes with relaxed selection in both specialists, exclusive of *D. gomerensis* and exclusive of *D. tilosensis*.

Supplementary Fig. 7. GO enrichment analysis of the genes with accelerated 1 kb upstream regions exclusive of *D. gomerensis* and exclusive of *D. tilosensis*.

